# Transcription Factor-Centric Approach to Identify Non-Recurring Putative Regulatory Drivers in Cancer

**DOI:** 10.1101/2022.01.31.478493

**Authors:** Jingkang Zhao, Vincentius Martin, Raluca Gordân

**Affiliations:** Center for Genomic and Computational Biology, Duke University, Durham, NC 27708, USA; Program in Computational Biology and Bioinformatics, Duke University, Durham, NC 27708, USA; Department of Computer Science, Duke University, Durham, NC 27708, USA; Department of Biostatistics and Bioinformatics, Duke University, Durham, NC 27708, USA; Department of Molecular Genetics and Microbiology, Duke University, Durham, NC 27708, USA

## Abstract

Recent efforts to sequence the genomes of thousands of matched normal-tumor samples have led to the identification of millions of somatic mutations, the majority of which are non-coding. Most of these mutations are believed to be passengers, but a small number of non-coding mutations could contribute to tumor initiation or progression, e.g. by leading to dysregulation of gene expression. Efforts to identify putative regulatory drivers rely primarily on information about the recurrence of mutations across tumor samples. However, in regulatory regions of the genome, individual mutations are rarely seen in more than one donor. Instead of using recurrence information, here we present a method to identify putative regulatory driver mutations based on the magnitude of their effects on transcription factor-DNA binding. For each gene, we integrate the effects of mutations across all its regulatory regions, and we ask whether these effects are larger than expected by chance, given the mutation spectra observed in regulatory DNA in the cohort of interest. We applied our approach to analyze mutations in a liver cancer data set with ample somatic mutation and gene expression data available. By combining the effects of mutations across all regulatory regions of each gene, we identified dozens of genes whose regulation in tumor cells is likely to be significantly perturbed by non-coding mutations. Overall, our results show that focusing on the functional effects of non-coding mutations, rather than their recurrence, has the potential to identify putative regulatory drivers and the genes they dysregulate in tumor cells.

## INTRODUCTION

Studies of somatic mutations in cancer genomes are generally focused on mutations that alter the amino acid sequences of protein-coding genes. However, whole-genome sequencing of human tumors has revealed that the vast majority of somatic mutations in cancer are non-coding (ICGC/TCGA Pan-Cancer Analysis of Whole Genomes Consortium 2020), suggesting that they could play a role in cancer initiation and development. Tumorigenesis is thought to be due to the accumulation of multiple driver mutations that confer a growth advantage to the tumor cells; some of these driver mutations may be non-coding (Khurana et al. 2016). But only a small proportion of the mutations present in cancer cells are drivers, so it is important to accurately identify them and distinguish them from the much larger number of passenger mutations (Elliott and Larsson 2021).

Given that driver mutations are expected to be under positive selection, their identification is generally based on patterns of recurrence among tumor samples. Several recent studies have attempted to discover non-coding driver mutations in regulatory DNA sites using recurrence information (e.g. Lochovsky et al. 2015, 2018; Rheinbay et al. 2017; Weinhold et al. 2014). Such studies usually involve the identification of genomic regions with high mutational frequency (i.e. hotspots) by comparing the mutation rate within a DNA window to a background distribution. However, it is generally challenging to precisely estimate the background mutation rate in small genomic regions, given the heterogeneity across different patients and across the genome (Lawrence et al. 2013). A recent meta-analysis of methods for predicting regulatory driver mutations reported that hotspot-based methods can generate large sets of candidate drivers, many of which are false positives (Rheinbay et al. 2020). To narrow down the list of candidates, one can also incorporate information on the functional impacts of putative non-coding driver mutations, in particular their effect on transcription factor (TF) binding. One of the most widely used approaches to prioritize mutations in regulatory regions involves the identification of TF binding sites created or disrupted by the mutations, which can be predicted using position weight matrices (PWMs) and motif prediction algorithms (Link et al. 2018; Shen et al. 2020). However, such methods are limited by the high false positive and false negative rates of binding site prediction algorithms.

In addition, Rheinbay et al. (Rheinbay et al. 2020) have recently reported that non-coding regulatory driver mutations are much less frequent than protein-coding drivers, with the only notable exception being driver mutations in the *TERT* gene promoter (Horn et al. 2013; Huang et al. 2013). Moreover, some non-coding drivers identified in previous studies were found to be the result of poorly-understood localized hypermutation processes such as mutations originating from differential DNA damage (Buisson et al. 2019) or differential DNA repair (Mas-Ponte and Supek 2020; Sabarinathan et al. 2016; Perera et al. 2016). On the other hand, recent studies of cancer drivers have shown that mutations do *not* have to be highly recurrent in order to be true drivers (Kim et al. 2016); in fact, even mutations that occur in individual tumor samples can drive tumorigenesis.

Here, we describe a new method for analyzing non-coding cancer mutations in regulatory genomic regions (i.e. promoters and enhancers) with the goal of prioritizing mutations based on their effects on TF binding. Unlike previous methods for prioritizing putative non-coding drivers, our method does not rely on the recurrence of mutations across tumor samples. Instead, we consider mutations to be potentially ‘significant’ if they lead to larger changes in TF binding affinity than expected by chance in that particular genomic region. Thus, the magnitude of the mutations’ effects, rather than their recurrence, is the basis for prioritizing mutations and regulatory regions for further studies.

To predict the quantitative effects of non-coding variants on TF binding we use QBiC-Pred (Martin et al. 2019), a computational method based on regression models of TF-DNA binding specificity trained on high-throughput *in vitro* data (Zhao et al. 2017). We focus on single-nucleotide mutations, since they are the dominant type of somatic mutation identified in cancer genomes (Khurana et al. 2016). Importantly, our method links enhancers and promoters to the genes they are likely to regulate, and it combines evidence from all regulatory regions of each gene in order to infer whether a gene is potentially dysregulated due to non-coding mutations. Finally, we use gene expression data from donors with versus without mutations in promoters and enhancers in order to validate that our method prioritizes biologically relevant mutations and regulatory regions.

## METHODS

### ICGC simple somatic mutations and gene expression data

To develop and test our new method for prioritizing putative regulatory driver mutations, we used the Liver Cancer-RIKEN, Japan (LIRI-JP) project from the International Cancer Genome Consortium (ICGC) (ICGC/TCGA Pan-Cancer Analysis of Whole Genomes Consortium 2020). We chose this project because it has a large number of donors with whole-genome simple somatic mutation (SSM) data (258 donors), as well as gene expression data (RNA-seq) for 230 out of the 258 donors with SSM data. The study reported a total of ∼3.8 million mutations, most of which are single nucleotide mutations (∼3.5 million). We used these mutations in our analyses.

### Promoter and enhancer data

We focused our analyses on mutations within promoter and enhancer regions, as TF binding sites are located in these regions. We defined promoters as the genomic sequences within +/- 1000 bp of each RefSeq (O’Leary et al. 2016) transcription start site (TSS), excluding any RefSeq exon sequences. We focused on promoters of protein-coding genes, using only TSSs that map to genes within the HUGO gene nomenclature (HGNC) (Tweedie et al. 2021). These criteria resulted in a set of 21,543 promoters.

For enhancers, we used the experimentally determined enhancers from the FANTOM5 project (Andersson et al. 2014; Lizio et al. 2015), which are frequently used in studies of non-coding mutations (e.g. Weinhold et al. 2014; Khurana et al. 2016). Importantly, the FANTOM consortium provides information about the linkage between enhancers and associated TSSs, which is critical for being able to connect our enhancer results to gene expression data. After removing enhancers that overlapped with promoter regions, we obtained a total of 41,254 enhancers.

We further filtered the 21,543 promoters and the 41,254 enhancers to keep only those that contain mutations. Since most enhancers are hundreds of base pairs long (median=254bp) and promoters are ∼2,000 bp, the majority of enhancers and a fair number of promoters do not contain any mutations (Figure S1). Therefore, after removing these regulatory regions without mutations, we obtained a set of 12,612 promoters and 9,018 enhancers with mutations in the LIRI-JP study.

### Defining the effects of mutations on TF binding, and the significance of these effects

We use TF binding changes to prioritize mutations that might act as drivers within each regulatory region (enhancer or promoter). For each region, we ask whether the mutations detected in that region lead to *larger* TF binding changes than expected by chance, i.e. according to a background model of random mutations (Figure 1A). To assess the effect of non-coding mutations on TF-DNA binding we use QBiC-Pred (Martin et al. 2019), a method we recently developed to quantify TF binding changes based on regression models trained on high-throughput *in vitro* binding data. While other binding specificity models can be used to predict the effects of mutations on TF binding, here we use QBiC-Pred because it performed better than methods based on position weight matrix or deep learning models of specificity (Martin et al. 2019; Zhao et al. 2017).

**Figure 1.**
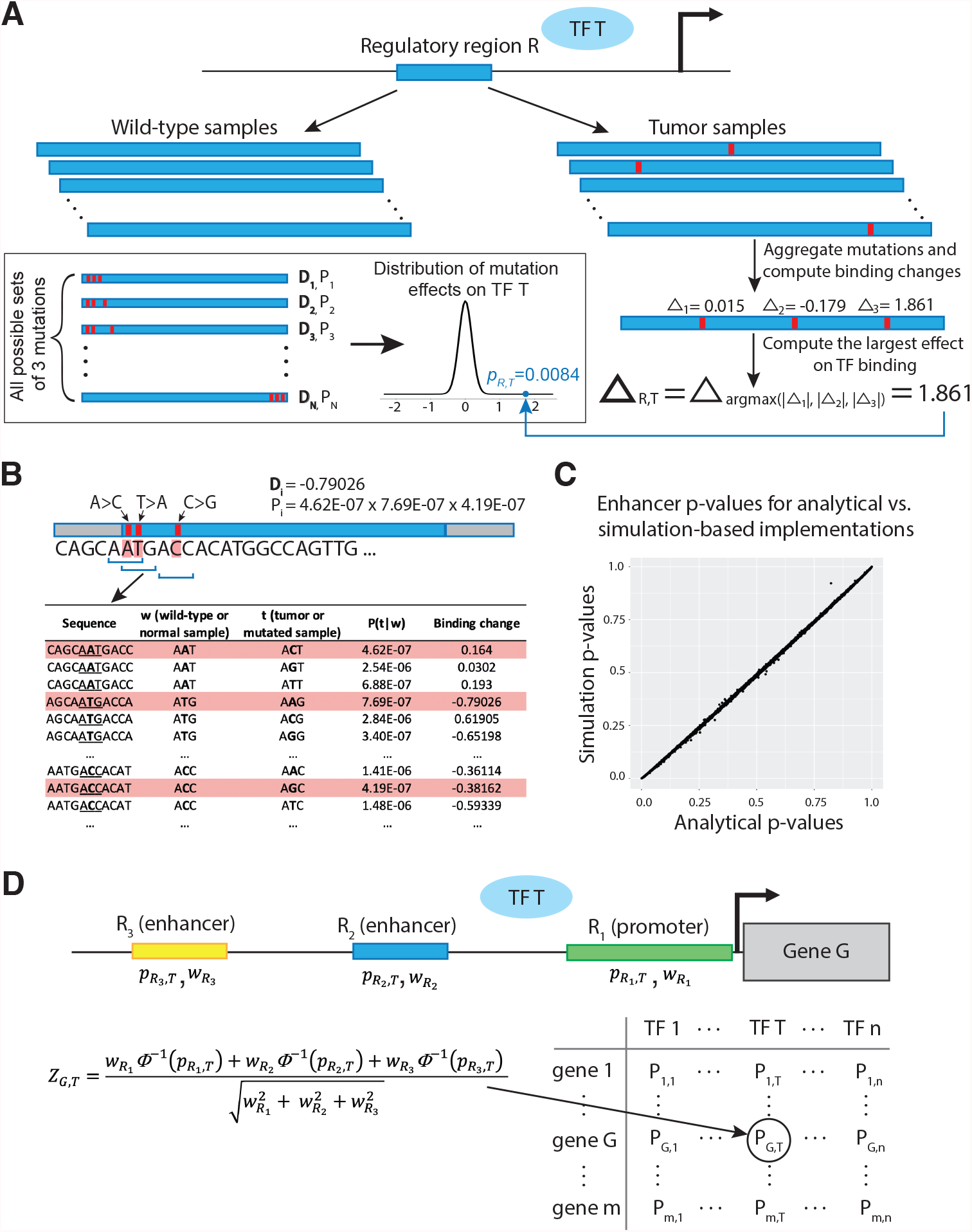
TF-centric approach to prioritize genes based on mutations in their regulatory regions. **(A)** For each regulatory region *R* and TF *T*, we aggregate the mutations across all patients in the cohort of interest, and compute the largest effect (either positive or negative) across all mutations. This effect, **Δ**_*R,T*_, is then compared to the distribution of effects computed for all possible sets of *k* mutations, where *k* is the number of actual mutations observed in region *R* (here, *k* = 3). **(B)** For any set *i* of *k* mutations, we compute the binding change with the largest magnitude among these *k* mutations (**D***i*), and the probability of that set of *k* mutations (*P*_*i*_), as described in Methods. **(C)** Comparison of p-values computed for mutation effects on MYC binding, for 9,018 enhancers. Plot shows the high correlation between p-values calculated using the analytical versus the simulation-based approach. **(D)** For genes with multiple regulatory regions, we compute the combined significance of the TF binding changes for TF *T* across all regions *R*_*i*_ by combining their p-values 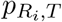 using Liptak’s method, also known as weighted Stouffer’s method, as described in Methods. We then use the combined p-values, adjusted for multiple hypothesis testing, to rank genes according to the smallest p-value across all TFs.

For each mutation of interest, QBiC-Pred reports its predicted effect on the binding specificity of 582 human TFs, based on models derived from 667 universal protein binding microarray (PBM) data sets. The effect of a mutation *m* on TF *T* is reported in terms of the difference (Δ_*m*_) in the logarithm of the PBM binding intensity signal for the mutated sequence relative to the wild-type sequence according to the binding model for TF *T*, as described in detail in (Martin et al. 2019). Positive values represent increased TF binding, while negative values represent decreased binding. Although here we focus on binding changes predicted with QBiC-Pred, our approach can directly use other binding specificity models, as long as they accurately reflect the quantitative TF binding changes induced by DNA mutations.

For a transcription factor *T* and a regulatory region *R* that has one or more mutations in the data set of interest, we compute the largest effect on TF binding (either positive or negative) over all mutations in *R* (**Δ**_*R,T*_, Figure 1A). Next, to determine if this effect is significant, we compare it against the distribution of effects expected by chance, according to a background model that takes into account: 1) the mutation spectra in that particular cohort, and 2) the particular DNA sequence of regulatory region *R*. Conceptually, if there are *k* total mutations in region *R* (*k* = 3 in Figure 1A), the full distribution of possible binding effects will be computed taking into account all possible sets of *k* mutations across the region. Each set *i* of *k* mutations will have a particular effect on TF binding (**D**_**i**_) and will occur with a particular probability (*P*_*i*_) (Figure 1B). The effect **D**_**i**_ is computed by taking the largest effect over the *k* mutations, similarly to the case of the real mutations. The probability *P*_*i*_ of a particular set of *k* mutations is computed as described below.

Using all single-nucleotide somatic mutations reported in the LIRI-JP study, we computed the mutation spectra for this cohort, analyzing mutations in their trinucleotide contexts (Alexandrov et al. 2020). As in previous studies (Alexandrov et al. 2020; Jusakul et al. 2017), we consider 6 mutation types (C>A,C>G,C>T,T>A,T>C,T>G) each in 16 possible contexts, based on the nucleotide right before and right after the mutated position. Mutations and their reverse complements (e.g. A**T**G>A**C**G and C**A**T>C**G**T) are counted together. For each trinucleotide, we estimate the probability of mutating the central base by taking the ratio between how many times that base was mutated in that context and how many times the wild-type trinucleotide occurs in regulatory regions (enhancers or promoters) across all normal samples. For example, the probability of mutating T in an A**T**G context in enhancers is:

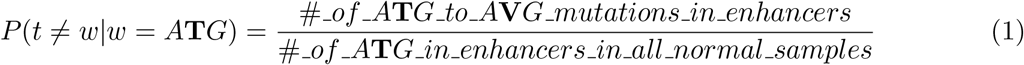

where *t* is the trinucleotide in the tumor sample, *w* is the trinucleotide in the corresponding normal sample, and V is the IUPAC code for {A, C, G} (i.e. not T). We calculate mutation probabilities separately for enhancers versus promoters, since they may be affected differently by mutagenic processes; indeed, we saw significant differences between the mutation spectra at enhancers versus promoters (Figure S2).

When the central nucleotide of a trimer is mutated, i.e. *t* ≠ *w*, there are three possible mutations, e.g. A**T**G to A**A**G, A**C**G, or A**G**G. We estimated the probability of each mutation type (e.g. A**T**G to A**A**G), given that a mutation exists at the central nucleotide, as the number of times we observed that particular mutation type divided by the total number of times the central nucleotide was mutated in that context, in the regulatory regions of interest. For example, focusing on enhancer regions, we compute:

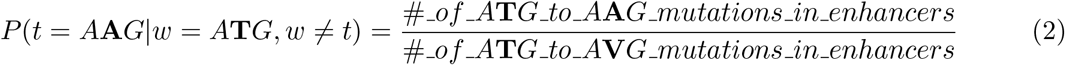

Next, to calculate the probability of a particular mutation in a particular trinucleotide context, we multiply the probability that the trinucleotide is mutated with the probability of the specific mutation, e.g.:

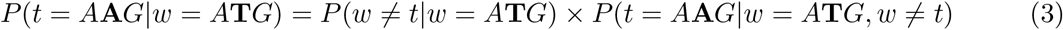

which can be simplified to the number of A**T**G to A**A**G mutations in enhancers divided by the total number of A**T**G trinucleotides in enhancers across all normal samples.

We note that the *k* mutations aggregated over region *R* are typically from different samples in our cohort, and can thus be considered independent. Therefore, for a set *i* of *k* mutations we compute the overall probability *P*_*i*_ of that particular set by multiplying the individual probabilities *P* (*t*|*w*) for each of the *k* mutations, as illustrated in Figure 1B. Finally, to assess the significance of **Δ**_*R,T*_, we compare this value against the distribution of effects for random sets of *k* mutations in region *R*, with the p-values being computed efficiently from *P*_*i*_ and **D**_*i*_ values using either an analytical or a simulation-based approach, as described below.

### Analytical and simulation-based approaches to compute the significance of mutation effects on TF binding

Our simulation-based approach uses the mutation probabilities described above to repeatedly sample mutations in the regulatory region of interest. Let us consider a regulatory region *R* of length *L* that contains *k* mutations across all patients in our cohort. Since there are 3 possible mutations for each position in *R*, we have a total of 3*L* mutations to consider in this region. At each iteration of our simulation-based approach, we randomly sample *k* out of the 3*L* possible mutations, with replacement. We do the sampling with replacement because the exact same mutation can occur in two or more patients. For this sampling process, the probability of selecting a particular set *i* of *k* mutations (*P*_*i*_) is computed by multiplying the probabilities of the *k* mutations (Figure 1B). The TF binding change for this set of mutations (**D**_*i*_) is computed by taking the maximum effect over the *k* randomly chosen mutations, as described above and illustrated in Figure 1A,B. By repeatedly sampling sets of *k* mutations using this procedure, we can approximate the distribution of mutation effects on TF binding (Figure 1A), and use it to compute empirical p-values for **Δ**_*R,T*_, taking the sign of the effect into account. This simulation-based approach is simple to understand and implement. However, simulations are time consuming and unfeasible for generating background distributions of mutations effects for all regulatory regions (totalling 12,612 promoters and 9,018 enhancers) and all TFs (totalling 582 TFs with 667 binding models available).

Alternatively, we can use an analytical approach to directly compute the p-value for the effect **Δ**_*R,T*_. Conceptually, the p-value of interest is the probability of obtaining an effect on TF binding at least as large as **Δ**_*R,T*_ when we randomly choose *k* of the 3*L* possible mutations in the regulatory region *R*. For simplicity, let us consider these effects in absolute value. For a set *i* of *k* mutations, if *at least one* of the mutations leads to an absolute change in binding of TF *T* that is ≥ |**Δ**_*R,T*_ |, then |**D**_*i*_| ≥ |**Δ**_*R,T*_ |. On the other hand, if *all* the *k* mutations lead to absolute binding changes < |**Δ**_*R,T*_ |, then we have |**D**_*i*_| < |**Δ**_*R,T*_ |. Thus, focusing on the absolute values of the effects of mutations on TF binding, we can compute our p-value of interest as:

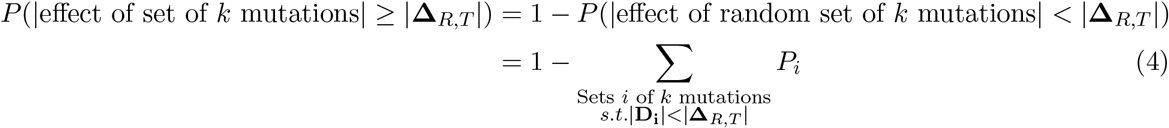

The total number of possible sets *i* of *k* mutations in regulatory region *R* is 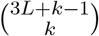, which is the number of possible unordered outcomes when sampling *k* out of 3*L* mutations with replacement. Even when choosing only the sets for which all *k* mutations have absolute binding changes < |**Δ**_*R,T*_ |, the number of possibilities can be very large and not feasible to compute explicitly. To overcome this problem, we compute a vector *π* = (*π*_1_, *π*_2_, …, *π*_*l*_), 0 ≤ *l* < 3*L*, with the probabilities of all individual mutations *m* in region *R* for which |Δ_*m*_| < |**Δ**_*R,T*_ |. The sum in Equation 4 can then be written in terms of the vector *π*, as the sum of the all elements in the outer product of *π* with itself, taken *k* times. For example, for *k* = 3, each element *P*_*i*_ in Equation 4 is an element of *π* ⊗ *π* ⊗ *π*. Importantly, we do not need to compute this outer product explicitly, as we are only interested in the sum of all elements in the product, which can be written as (*π*_1_ + *π*_2_ + … + *π*_*l*_)^*k*^. Thus, our p-value of interest can be calculated as:

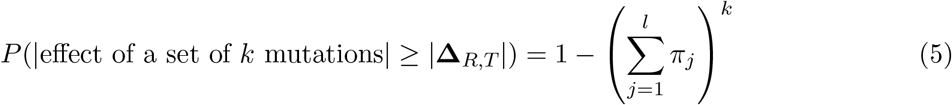

Finally, since we want to take into account the directionality of the effect of mutations on TF binding, i.e. decreased binding (**Δ**_*R,T*_ < 0) or increased binding (**Δ**_*R,T*_ > 0), we calculate the one-sided p-value as 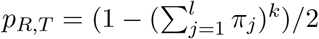.

We confirmed that the results from our analytical and simulation-based approaches agree with each other (Figure 1C). For this test, we randomly picked one of the TFs, MYC, and we calculated the p-values for changes in MYC binding specificity for all 9018 mutated enhancers, using a simulation process with one million iterations. Since the simulation-based approach is more time consuming and has limited precision in estimating the p-values of interest (due to the limited number of iterations), we used the analytical approach for all subsequent analyses.

### Integrating results across all regulatory regions of a gene

Genes encoded in the human genome typically have multiple regulatory regions (enhancers and promoters); mutations in either of these regions could affect a gene’s expression. Thus, it is of interest to integrate the effects of mutations across all regulatory regions of each gene. As detailed above, we define gene promoters based on TSS coordinates in the RefSeq database (O’Leary et al. 2016), and we leverage TSS-enhancer links from the FANTOM consortium (Lizio et al. 2015), considering all the cell types and tissues with available data. In other words, if a genomic region has been identified as an enhancer for gene *G* in one tissue, then we consider that region as part of the regulatory landscape of gene *G*, in order to be as inclusive as possible. On average, each TSS is associated with 4.9 enhancers according to the FANTOM data. For genes that have multiple TSSs in RefSeq, we consider each TSS separately. Thus, some genes may appear multiple times in our final results, with different p-values that correspond to its different TSSs.

Given a transcription factor *T* and a gene *G* with *r* regulatory regions containing at least one mutation in our cohort of interest, we calculate the significance 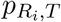 of the effects 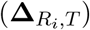 of mutations in each regulatory region *R*_*i*_ on the binding specificity of TF *T*. Next, we want to integrate these effects over all *r* regulatory regions by combining their p-values. Intuitively, our null hypothesis (*H*_0_) is that the effects for all regulatory regions 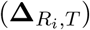 come simply from the background distribution of effects due to random mutations. The alternative hypothesis (*H*_1_) is that at least one regulatory region of gene *G* has an effect 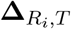 significantly larger than expected by chance according to the background model of mutations in regulatory regions. Importantly, as the promoter and enhancer regions used in our analysis do not overlap, we can consider the p-values 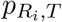 as coming from independent tests.

One approach to combine the *r* p-values computed for gene *G* is Fisher’s method (Fisher 1934), which is often used in meta-analyses, including analyses of non-coding mutations in cancer (Rheinbay et al. 2020; Lawrence et al. 2014; Araya et al. 2016). However, Fisher’s method would not take into account the fact that different regulatory regions have different lengths and different probabilities of harboring mutations. If a regulatory region *R*_*i*_ is very long, then the number of possible mutations, and thus the number of possible effects on TF *T*, is also large. In comparison, a shorter regulatory region, *R*_*j*_, will have fewer possible mutations and fewer possible effects on TF *T*. If the two regions have the same p-value, i.e. 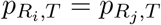, we might still consider, intuitively, that the mutation effect in *R*_*i*_ is more significant because for *R*_*i*_ it would be easier to achieve a large effect on TF binding, since more mutations can occur in *R*_*i*_ than *R*_*j*_. This is similar to meta-analyses of studies with very different sample sizes, where a weighted version of Stouffer’s method, developed by Liptak (Lipták 1958) and also known as the weighted Z-method or weighted Z-test, was found to be superior to Fisher’s method when combining p-values from independent tests (Whitlock 2005; Zaykin 2011).

Here, we use Liptak’s method (Lipták 1958) to combine the p-values of all regulatory regions of a gene, with the weights computed based on the mutations probabilities in each region, combined using Shannon’s entropy. Specifically, for a regulatory region *R* of length *L* we compute its weight as 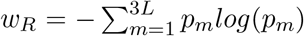, where *p*_*m*_ is the probability of the *m*^*th*^ possible mutation, computed according to the trinucleotide centered at that position (see Figure 1B). We note that weights computed in this manner are overall correlated with the length of the regulatory regions, but avoid giving an out-sized importance to very long regions (Figure S3). Thus, for TF *T* and gene *G* with *r* regulatory regions, we compute the weighted test statistic 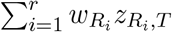, where 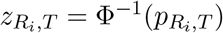 and Φ^−1^ is the inverse of the standard normal cumulative distribution function, as initially proposed by Liptak (Lipták 1958; van Zwet and Oosterhoff 1967). Under the null hypothesis (*H*_0_), this test statistic follows a normal distribution 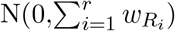 for any choice of weights (van Zwet and Oosterhoff 1967; Heard and Rubin-Delanchy 2018). This allows us to compute the p-value of the combined test, *P*_*G,T*_, as follows:

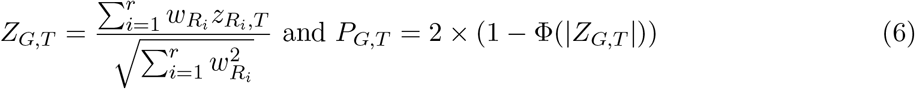

where Φ is standard normal cumulative distribution function. Finally, we used the Benjamini-Hochberg correction (Hochberg 1988) to adjust for multiple hypothesis testing across all genes and all TFs. We then ranked genes according to the smallest p-value across all TFs, and we analyzed the top genes for differences in gene expression. In total, we analyzed 5,336 genes with mutations in enhancers, 11,721 genes with mutations in promoters, and 13,982 genes with mutations in at least one regulatory region (enhancer or promoter).

## RESULTS

### Integrated analysis across regulatory regions identifies 54 genes with significant TF binding changes due to mutations in regulatory DNA

Using the single nucleotide mutation data from the LIRI-JP study, we identified 13,982 genes with mutations in either the promoter or the enhancer regions. For each of these genes, we analyzed all 582 human TFs for which QBiC-Pred models are available (Martin et al. 2019), and we computed the smallest p-value *P*_*G,T*_ across all TFs, adjusted for multiple testing, as described in the Methods section (Figure 1). We chose to take the minimum p-value, rather than integrate the p-values across factors, because TF binding specificities are oftentimes highly correlated, especially for closely-related paralogous TFs. Next, we ranked genes according to the smallest adjusted p-value across all TFs, and we identified 54 genes for which this p-value was < 0.05 (Figure 2, Table S1). In other words, for each of these 54 genes, our analysis revealed at least one TF for which the mutations in the regulatory regions of the gene have larger effects on TF binding specificity than expected by chance based on the mutation spectra in our cohort.

**Figure 2.**
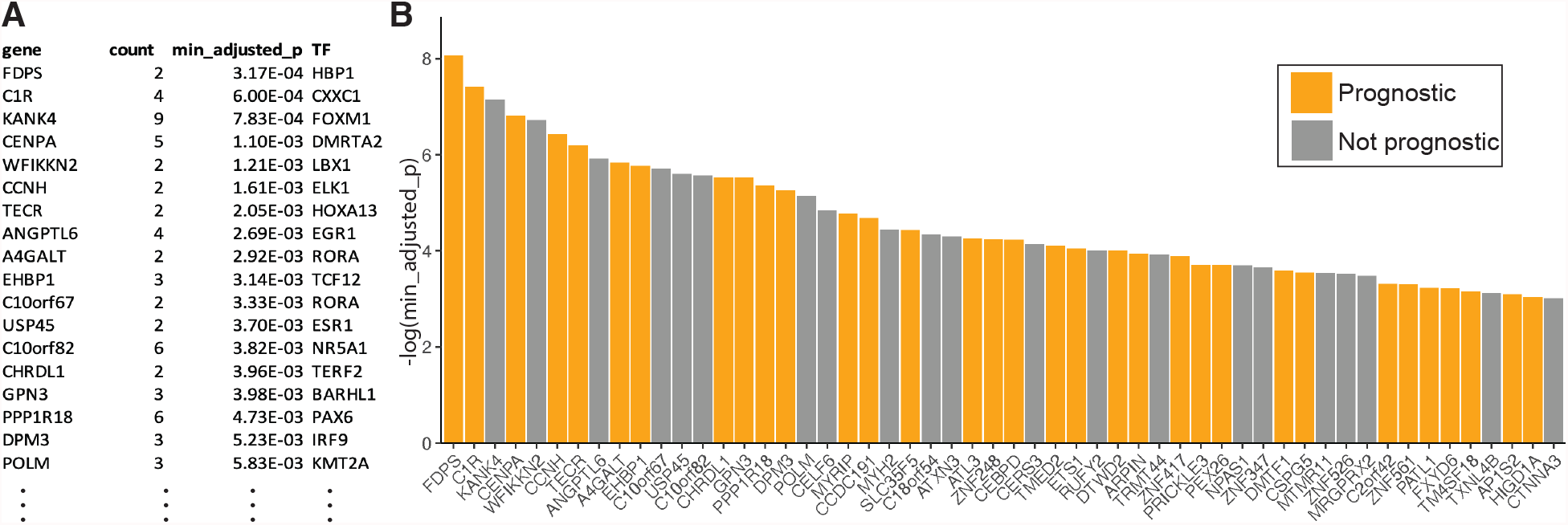
Genes prioritized based on mutations in their regulatory regions. **(A)** Top genes with the smallest adjusted p-values. The table shows each gene’s name, the number of mutations in all its regulatory regions, the minimum p-value after adjusting for multiple testing, and the TF with the most significant binding changes for that gene. **(B)** Barplot showing the top 54 prioritized genes from our combined promoter and enhancer analysis. Y-axis shows the negative logarithm of the minimum adjusted p-value. Cancer prognostic markers, as reported in The Human Protein Atlas, are shown in yellow.

To determine if these 54 prioritized genes are potentially relevant in cancer, we first asked whether this set is enriched for cancer prognostic genes. Using pathology data from The Human Protein Atlas (Uhlen et al. 2017), we found that 33 out of the 54 prioritized genes are indeed prognostic markers in at least one cancer type (Figure 2B). This represents a significant enrichment (*p* = 0.095, Fisher’s exact test) when compared to genes ranked in the bottom half of our ranked list. In addition, the top gene on our prioritized list (FDPS), as well as four other genes on the list (CENPA, PEX26, FXYD6, and TM4SF18) are prognostic markers in liver cancer.

We performed similar analyses focusing only on promoters or only on enhancer regions. For the enhancer-only analysis, we identified 5 significant genes (at a minimum adjusted p-value cutoff of 0.05) among the 5,336 genes with enhancer mutations (Table S1). For the promoter-only analysis, we identified 73 significant genes among the 11,721 genes with promoter mutations (Table S1), none of which were also prioritized in the enhancer-only analysis. These results are not surprising, given that some genes only have either enhancer or promoter mutations, but not both. In addition, here we use a stringent set of enhancers, as reported by the FANTOM consortium, in order to limit the number of false positive enhancer calls; however, there are likely a large number of false negatives, i.e. enhancers that are missing from our data. Among the 54 genes identified in the combined analysis of promoters and enhancers, 4 of them are also prioritized in the enhancer-only analysis, and 47 of them are prioritized in the promoter-only analysis (Figure S4). Three genes, ETS1, CELF6, and PALT1 were identified only in the combined analysis (Figure 2B).

We also found a significant enrichment of cancer prognostic genes in the set of 73 genes prioritized in the promoter-only analysis (44 of the 73 prioritized genes are prognostic markers, Fisher’s exact test *p* = 0.084). For the enhancer-only analysis, we found that 3 of the 5 prioritized genes are prognostic markers. However, given the small number of genes prioritized in this analysis, the enrichment in prognostic markers was not significant.

### Genes with significant mutations in their regulatory regions show large expression differences in mutated versus non-mutated samples

For the 54 genes prioritized based on mutations in enhancers and promoters, we also asked whether the mutations are likely to affect gene expression. To test this, we leveraged the gene expression data (EXP-S) available in ICGC for our cohort of interest, LIRI-JP (Methods). For each gene, we compared its expression level (i.e. normalized read counts, or normalized TPM values) for donors with versus without mutations in the regulatory regions of that gene, and we used a Wilcoxon rank-sum test to assess the significance of the observed gene expression differences. Our analysis revealed that the difference in gene expression between donors with vs. without mutations in regulatory regions are much more significant for the set of 54 prioritized gene (i.e. those with minimum adjusted Liptak’s test *p* < 0.05) compared to a control set of genes (i.e. those with *p* ≥ 0.1) (Figure 3A). Gene with intermediate p-values (0.05 ≤ *p* < 0.1) also showed significant gene expression differences, although less so than the top 54 prioritized genes (Figure 3A).

**Figure 3.**
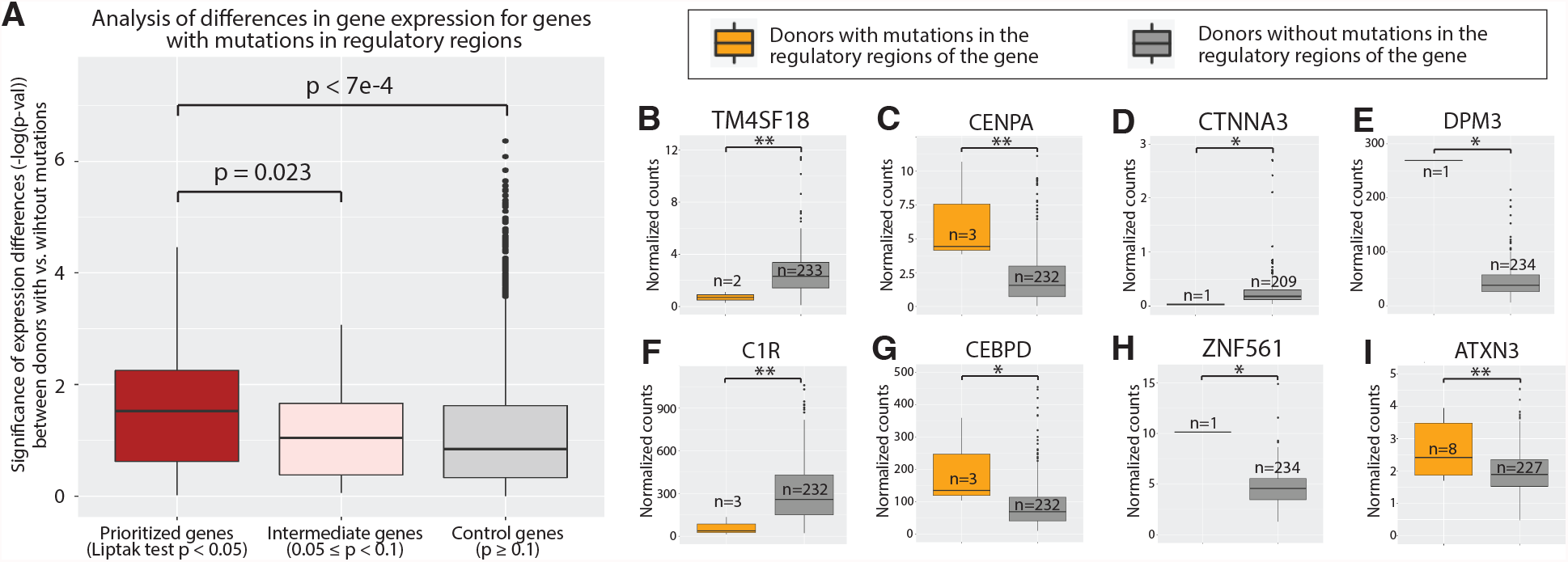
Analysis of differences in gene expression for genes with mutations in enhancer and promoter regions, comparing donors with vs. without mutations in regulatory DNA. **(A)** Genes prioritized by our analysis (minimum adjusted Liptak’s test *p* < 0.05) have larger expression differences due to mutations in regulatory regions compared to intermediate (0.05 ≤ *p* ≤ 0.1) and control (*p* ≥ 0.1) genes. To represent the expression differences (y-axis), we show the negative logarithm of the Wilcoxon rank-sum test p-values between the gene’s expression level in the two donor groups, i.e. with vs. without mutations in regulatory regions. **(B-H)** Boxplots showing the 8 prioritized genes with significant gene expression differences. The y-axes show the gene expression levels, as normalized read counts, as reported in ICGC. P-values are calculated using a two-sided Wilcoxon rank-sum test, with **\*\*** denoting *p* < 0.05 and **\*** denoting 0.05 ≤ *p* < 0.1.

We note that gene expression analyses could not be performed for all genes, as for some genes with mutations in regulatory regions there was no expression data available from the donors where the mutations were observed. Among the 54 prioritized genes, 43 genes had expression data for one or more donors with mutations in enhancers or promoters, and in most of those cases the number of such donors was one, making it difficult to reach statistical significance. Nevertheless, we found significant gene expression differences for 8 of the 43 prioritized genes with available data: TM4SF18, CENPA, CTNNA3, DPM3, C1R, CEBPD, ZNF561 and ATXN3 (Figure 3B-H).

Among these genes, TM4SF18 and CENPA (Figure 3B,C) are prognostic markers in liver cancers according to The Human Protein Atlas (Uhlen et al. 2017). In addition, CENPA has been shown to be aberrantly expressed in hepatocellular carcinomas (HCCs) compared to non-tumor tissues (Li et al. 2011). Another one of the significant genes, CTNNA3, is a tumor suppressor in HCCs (He et al. 2016); according to our analysis, its expression is very low in the only donor with mutations in the regulatory regions of CTNNA3 (Figure 3D), which is consistent with the gene’s role as a tumor suppressor. Gene DPM3 is part of the DPM family, whose members were found to be significantly correlated with shorter overall survival in liver cancer patients (Li et al. 2020); in our analysis, the only one donor with mutations in the regulatory regions of DPM3 had a very high DPM3 expression level (Figure 3E). Genes C1R, CEBPD, and ZNF561 (Figure 3F,G,H) are prognostic markers in renal cancers, which have been shown to metastasize to liver (Bianchi et al. 2012).

The remaining prioritized gene with significant expression differences, ATXN3 (Figure 3I), was interestingly not among the genes characterized as prognostic markers in cancer. ATXN3 is a member of the deubiquitinating enzymes family (DUBs). These enzymes catalyze the removal of ubiquitin from protein substrates and regulate several aspects of protein fate. ATXN3 was found to play important roles in several tumours, for example by deubiquitinating PTEN in lung cancer (Sacco et al. 2014) and KLF4 in breast cancer (Zou et al. 2019), although a role for ATXN3 in liver cancer has not yet been reported. According to our analysis, mutations in the ATXN3 enhancer regions result in significant gain-of-binding mutations for TF RUNX1 (Figure 4), a protein involved in tumour initiation and development in hematopoietic cells and in several tissues (Taniuchi et al. 2012; Otálora-Otálora et al. 2019; Lie-A-Ling et al. 2020). Among the eight donors with mutations in ATXN3 enhancers, five have mutations that lead to significant changes in RUNX1 binding specificity, and for these five donors the expression level of ATXN3 is significantly higher than for all other donors (*p* = 0.011, Wilcoxon rank-sum test), despite the small sample size. Overall, these results show that our method is able to identify and prioritize relevant genes that are likely to be significantly affected by mutations in their regulatory regions.

**Figure 4.**
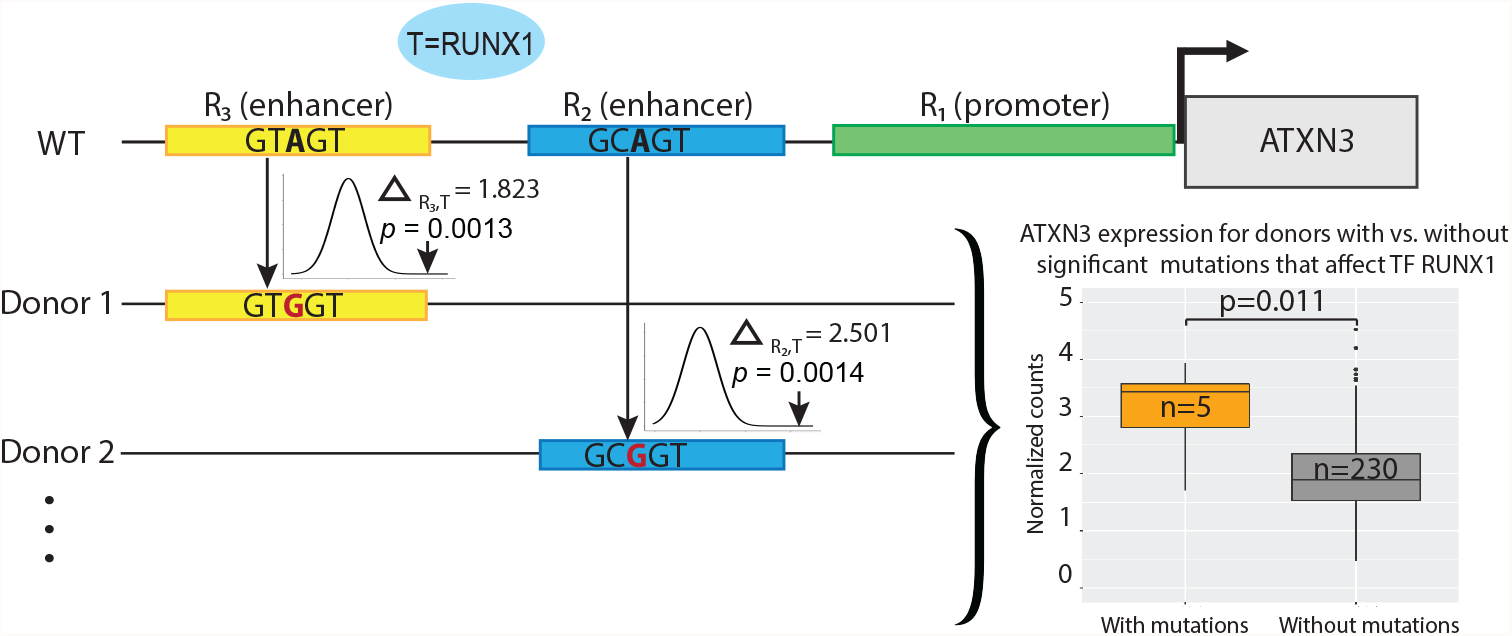
Mutations in the ATXN3 enhancers that alter the binding specificity of TF RUNX1. Out of 8 patients with ATXN3 regulatory mutations, 5 patients have significant (*p* < 0.1, one-sided Wilcoxon rank-sum test) mutations in enhancers which result in the gain of binding of TF RUNX1. Two enhancers where RUNX1 have the most significant binding changes among all TFs are shown. Donors with significant gain-of-binding RUNX1 mutations (orange boxplot) have higher ATXN3 expression compared to donors without ATXN3 regulatory mutations (grey boxplot), *p* < 0.011 two-sided Wilcoxon rank-sum test.

We also analyzed the expression levels of genes prioritized in the promoter-only and enhancer-only analyses. For the 5 genes prioritized according to mutations in enhancer regions, we found more significant expression differences (between donors with vs. without regulatory mutations) than in the control gene set (*p* = 0.03, one-sided Wilcoxon rank-sum test; Figure S6B). Interestingly, the set of 73 genes prioritized in the promoter-only analysis (Figure S5) did not differ significantly from a control set of genes (*p* = 0.231; Figure S6A). Of these 73 genes, 47 were also prioritized in the combined analysis of promoters and enhancers, while the remaining 26 genes appeared only in the promoter-only analysis, suggesting that the promoter-only analysis may be more prone to false positive, i.e. prone to prioritizing genes for which promoter mutations do not correspond to changes in gene expression. To further investigate this, we focused on the 26 genes, and found that only one of these genes (DDX21) has significant expression differences between donors with vs. without mutations (Figure S6C). This suggests that integrating regulatory mutations over enhancers and promoters, rather than promoters alone, has the best potential to prioritize genes that are dysregulated in cancer cells, likely due to non-coding mutations in the regulatory regions of the genes.

## DISCUSSION

In summary, we developed a new approach to identify putative regulatory driver mutations in cancer, based on quantitative predictions of the effects of single nucleotide variants on TF binding (Martin et al. 2019). Our method is orthogonal to existing tools (e.g. (Lochovsky et al. 2015, 2018; Rheinbay et al. 2017; Weinhold et al. 2014)), in that it does not require the putative driver mutations to be highly recurrent; instead, we assess the significance of the mutations by testing whether they cause *larger* TF binding changes than expected in the case of completely random mutations. Using this approach, we identified 54 potentially dysregulated genes (Figure 2) by prioritizing genes for which mutations in their enhancers or promoters lead to significant changes in TF binding specificity. Our analyses show that these genes are enriched for cancer prognostic markers, and they have higher differences in expression levels between donors with vs. without mutations in regulatory regions, compared to control gene sets (Figure 3). We also linked the potentially dysregulated genes with the TFs whose binding events are altered the most by putative regulatory mutations, such as shown in the example of ATXN3 (Figure 4).

We note that our method can be applied to any somatic mutation data. Here, we used single nucleotide mutations; however, our method can also be applied, with minor modifications, to small indels. Furthermore, we expect the results of our method to improve as higher quality data is used as input, including expanded sets of accurate TF-DNA specificity models, additional enhancer regions and accurate enhancer-promoter mappings. In addition, our method can be adapted for patient-level analyses. We note that current driver identification approaches combine mutations from different patients to gain more statistical power to identify significant regulatory mutations. However, a potential disadvantage of such approaches is that they may not work well on small cohorts or on cohorts that are highly heterogeneous. For approaches that identify drivers based on recurrence, it is impossible to run the analysis for individual patients. However, since our method does not require the driver mutations to be recurrent, it can be applied to identify potentially dysregulated genes for each patient, using our simulation-based approach, with minor modifications to the resampling process. Briefly, instead of using sampling with replacement, we would sample without replacement for patient-level analyses because a patient cannot have the same mutation more than once. The patient-level analysis would need to be further refined so that it has more coherent evidence to prioritize genes for follow-up experimental validations; however, it would provide a very different perspective than the cohort-level approaches.

Overall, the TF-centric approach described here uses a distinctive pipeline to identify putative regulatory driver mutations in cancer, by focusing on the magnitude of the effect of the mutations. Our approach is orthogonal to existing methods and thus serves to complement existing tools and resources for analyzing and prioritizing putative non-coding drivers (e.g. (Liu et al. 2021; Zhu et al. 2020; Zhang et al. 2018)). Our results show that most of the potentially dysregulated genes prioritized by our method either have large expression differences in donors with vs. without mutations, or are cancer prognostic genes, or both. Experimental validations (such as tumorigenicity experiments (Kim et al. 2016)) are needed to determine whether these genes actually contribute to cancer development. Nevertheless, our results suggest that regulatory mutations should be investigated further, not just based on their recurrence, but also based on their functional effects, in order to uncover dysregulated genes that may drive tumorigenesis.

## DATA ACCESS

The R code used in this work is available in GitHub at https://github.com/jz132/cancer-mutations.

