## Supplemental Figures for "Transcription Factor-Centric Approach to Identify Non-Recurring Putative Regulatory Drivers in Cancer"

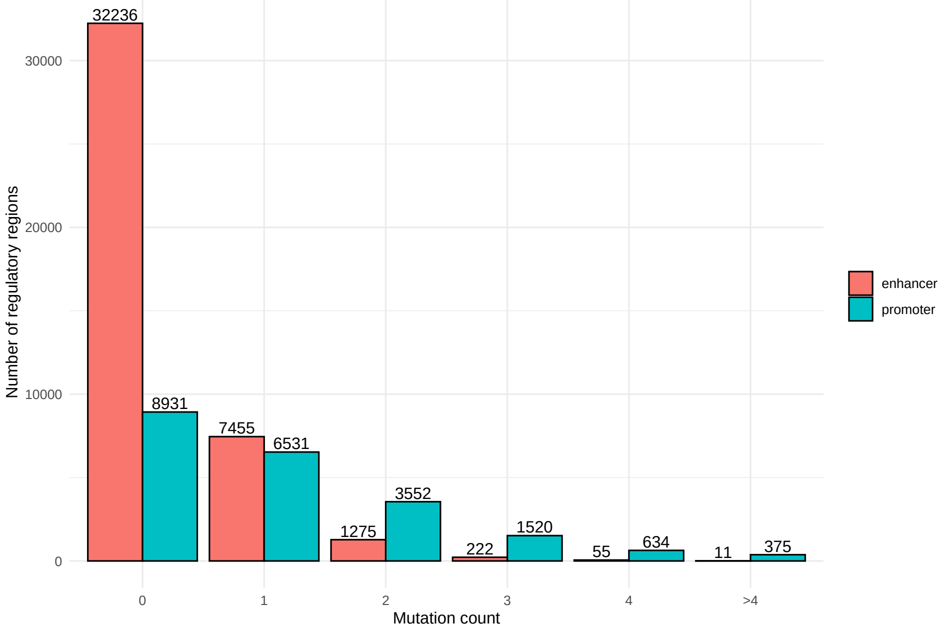


**Figure S1.** Number of mutations in regulatory regions. The average mutation rate is $3.6 \times 10^{-6}$ in enhancers and $2.1 \times 10^{-6}$ in promoters.

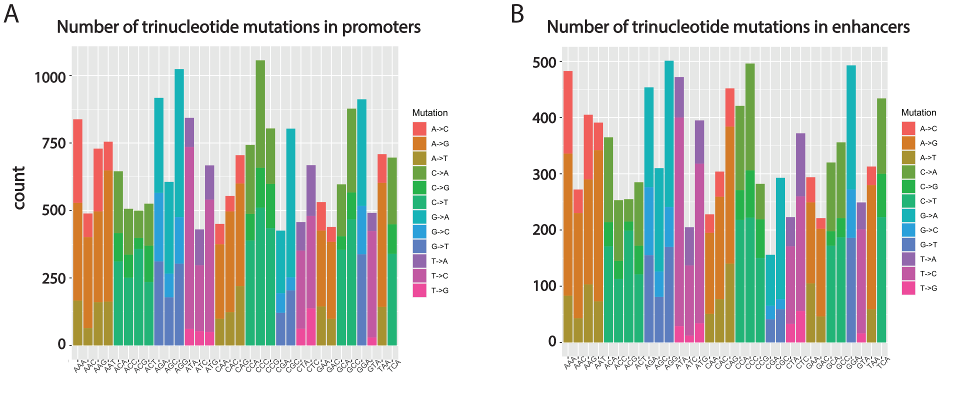


**Figure S2.** Number of mutations in promoters (A) and enhancers (B), computed for each mutation type in each trinucleotide context.

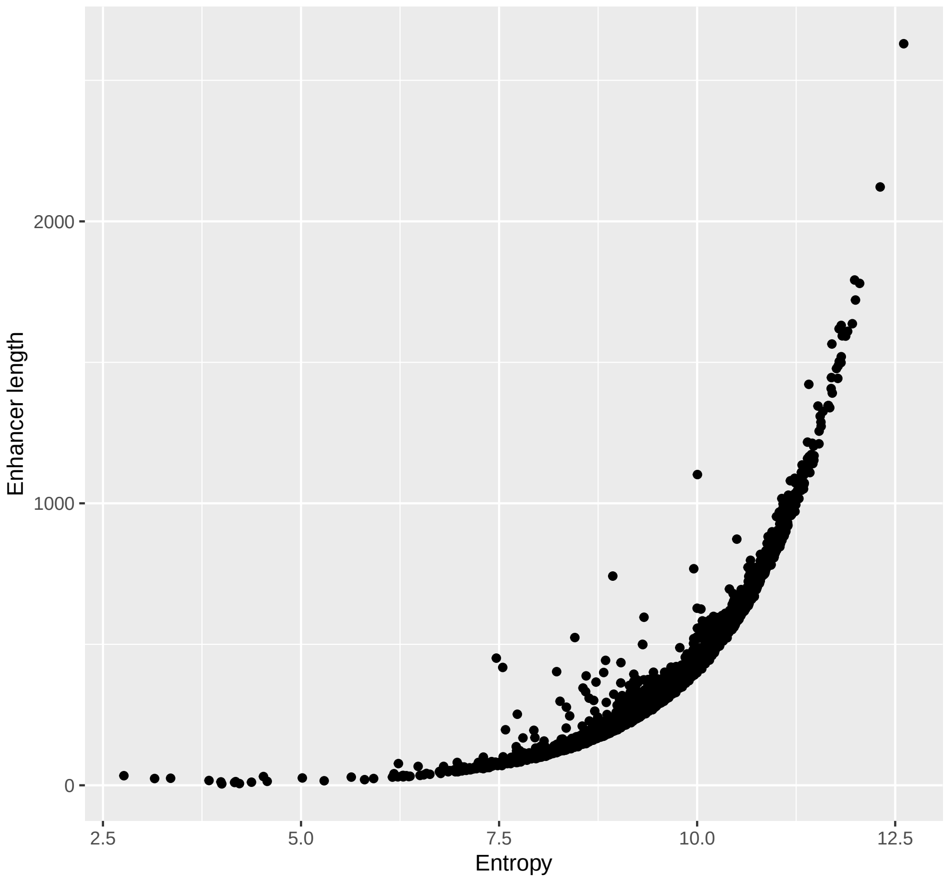


**Figure S3.** Comparison between the entropy computed for each enhancer, based on the probabilities of all possible mutations, versus the enhancer's length.

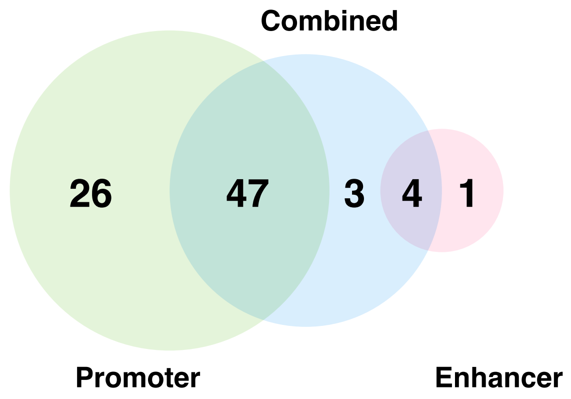


**Figure S4.** Number of prioritized genes in the promoter-only (green), combined (blue), and enhancer-only (pink) analyses.

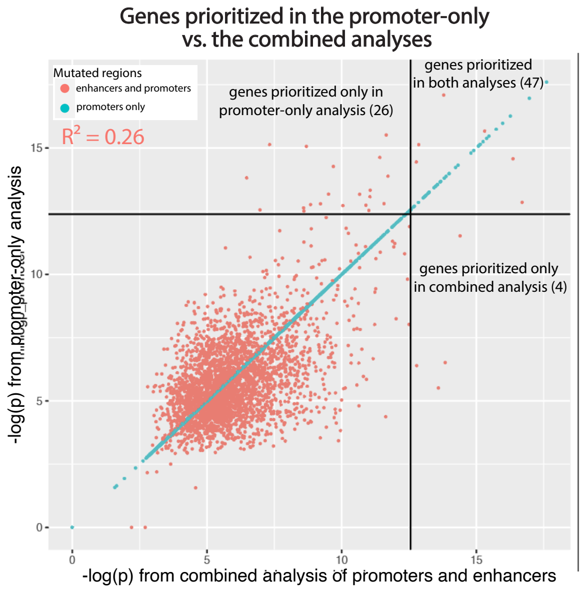


**Figure S5.** Comparison between the genes' significance in the promoter-only versus the combined analysis, shown as negative logarithm of the p-value. The $R^2$ shows the correlation for genes with mutations in both promoters and enhancers (shown in orange). Only genes with at least one promoter mutation were considered in this analysis.

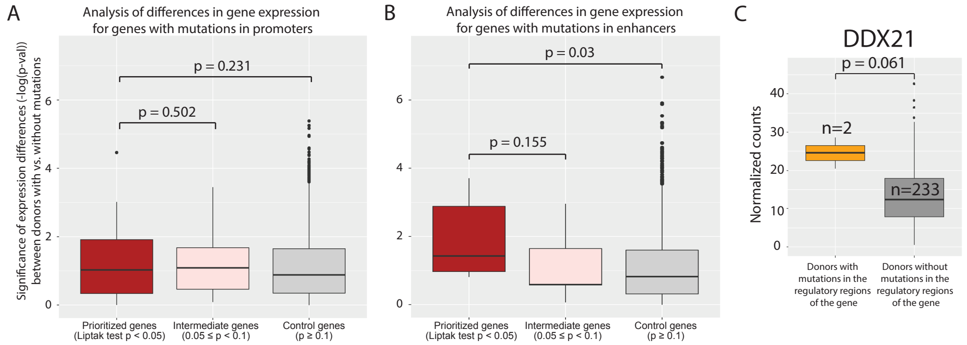


**Figure S6.** (A) Significance of gene expression differences between donors with vs. without mutations in promoters, shown for prioritized genes (minimum adjusted Liptak's test $p < 0.05$), intermediate genes ($0.05 \leq p < 0.1$) and control genes ($p \geq 0.1$) genes. (B) Similar to panel A, but analyzing mutations only in enhancers. (C) Only one gene (DDX21) that was prioritized in the promoter-only analysis but not in the combined analysis, has a significant expression difference between donors with vs. without mutations.
